# CORE: Deterministic alignment for bias-free organoid histology

**DOI:** 10.64898/2026.09.20.752946

**Authors:** Iago Pereiro, Barbora Lavickova, Capucine Labat-Berthier, Simone Toscani, Julien Aubert, Irineja Cubela, Jyoti Rao, Jose L Garcia-Cordero

## Abstract

Histological analysis of 3D tissue models is limited by geometric sampling bias arising from inconsistent positioning during embedding. We present Coplanar Organoid Reproducible Embedding (CORE), which aligns heterogeneous specimens on a shared sectioning plane to enable reproducible, bias-free histology. By eliminating depth-dependent sampling artifacts, CORE reveals biological differences obscured by sectioning depth and integrates with automated liquid-handling workflows.

## Main text

Spatial profiling of 3D tissue models is constrained by embedding methods that lack control over the axial position of individual specimens within the block^1–5^(**Figure 1a**). Therefore, a single microtome section rarely intersects the equatorial plane of all specimens simultaneously; in any given cut, some organoids are sectioned off-center while others are missed entirely^4,6^ (**Figure 1b**, empty wells). Even when specimens are captured, non-equivalent anatomical regions are compared across a cohort, introducing a geometric sampling bias that distorts measurements of tissue architecture and cellular composition. This bias can mask biologically meaningful differences between experimental conditions. Recovering true equatorial planes through serial sectioning is labor intensive^7^, while digital reconstruction from full section stacks substantially increases imaging, computational, and storage requirements^8^ for sections that are ultimately discarded. Critically, both approaches rely on comparing specimens across different physical sections, susceptible to inter-section variability in signal intensity, from assay processing to image acquisition, that confounds quantitative comparisons between individual specimens^9,10^. Existing active^4,6,11,12^ and passive^1,2,13–17^ alternatives partially mitigate this problem but remain susceptible to specimen heterogeneity and positional drift during embedding, resulting in persistent depth variability.

**Figure 1.**
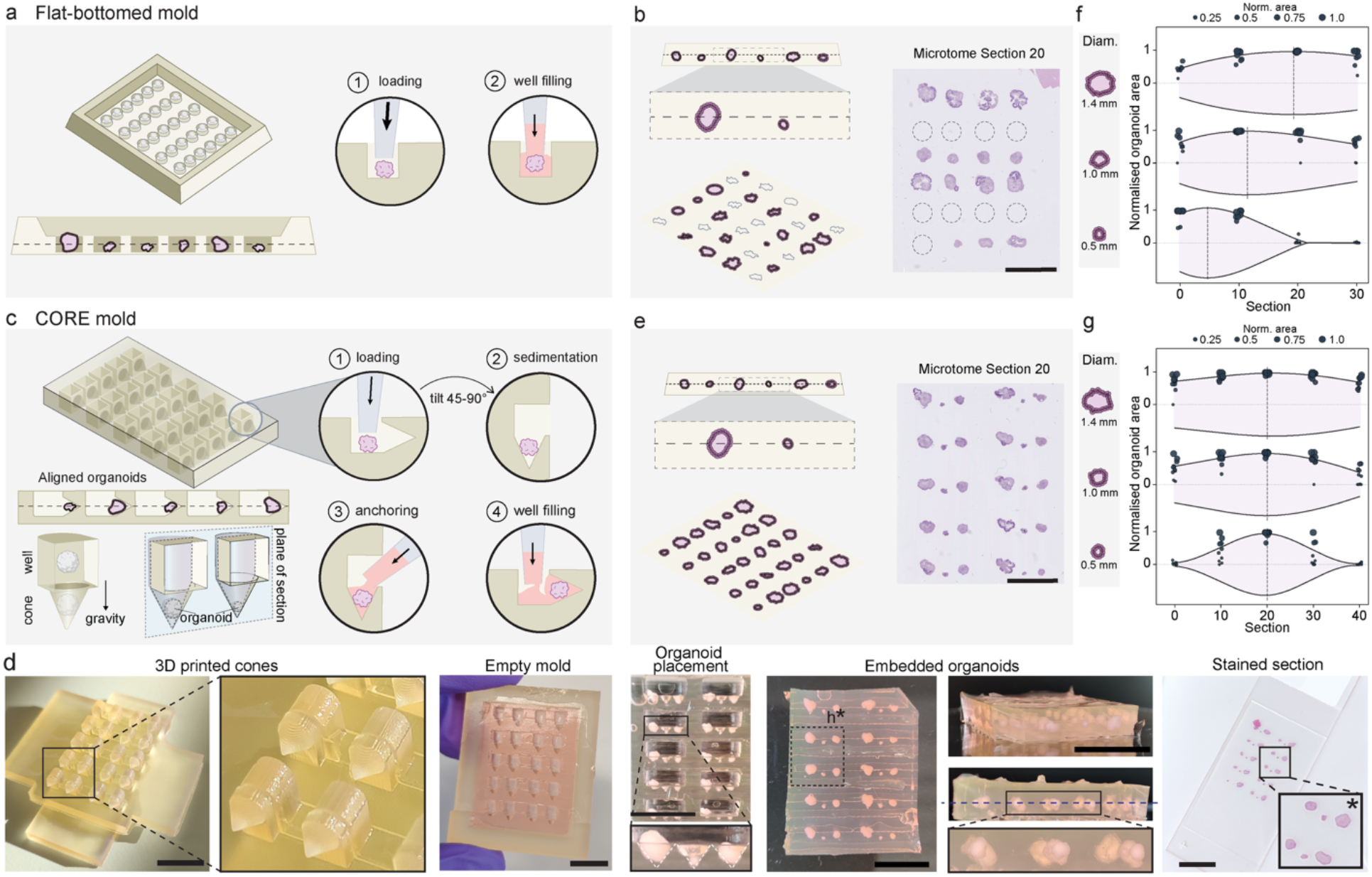
CORE eliminates depth-dependent positioning variability in heterogeneous organoid cohorts. **a**, Conventional flat-bottomed embedding, where size variation causes staggered depths and sampling errors. **b**, H&E section of varied-size retinal organoids in a flat-bottomed mold (dashed circles: empty wells). Scale bar, 5 mm. **c**, CORE platform schematic showing how gravity-driven conical alignment and apex anchoring achieves single-plane core capture. **d**, Workflow photographs: 3D-printed insert (cone-tip rounding is an optical artifact), cast hydrogel base, loaded 6×5 array, decasted gel showing coplanarity, and final H&E slide. Scale bars, 10 mm. **e**, H&E section of varied-size retinal organoids in a CORE mold. Scale bar, 5 mm. Equatorial-section depth versus normalized organoid size for flat-bottomed (**f)** and CORE (**g)** blocks demonstrating size-dependent depth variability in conventional embedding and convergence onto a shared sectioning plane with CORE (dashed lines shows size means).

To address this, we developed Coplanar Organoid Reproducible Embedding (CORE), which decouples specimen positioning from hydrogel gelation through passive, gravity-assisted sedimentation in pre-cast wells, without external positioning tools or manual orientation (**Figure 1c, Supplementary Figure 1**). The CORE platform consists of an array of L-shaped hydrogel wells cast from 3D-printed master molds (**Figure 1d, Supplementary Figure 2a-b**). Each well comprises a vertical loading chamber and a lateral conical apex perpendicular to it. Embedding proceeds in four stages: specimens are loaded into the wells; the mold is tilted to 45°-90° so gravity funnels each specimen into the apex; a localized anchoring hydrogel immobilizes them; the well is filled to produce a sectionable block (**Supplementary Figure 2c, Supplementary Video 1**). Because alignment is established before gelation, the geometric centers of heterogeneous specimens converge onto a shared sectioning plane largely independent of specimen size (**Figure 1d, e**), eliminating the depth variability that underlies geometric sampling bias.

We benchmarked CORE on heterogeneous human retinal organoids (0.5-1.4 mm). In flat-bottomed molds, the depth of the equatorial plane scaled with organoid diameter, with the three size classes centered at sections 5, 12, and 20, spanning up to 15 sections across the cohort (**Figure 1f, Supplementary Figure 2d**). Consequently, no single section captured all organoids, and quantitative comparisons were derived from non-equivalent anatomical planes. In contrast, CORE collapsed all three size classes onto a shared equatorial plane (**Figure 1g, Supplementary Figure 2e**), reducing axial spread to 2 sections, a 7.5-fold improvement, and effectively eliminating size-dependent variation in sampling depth.

Because large organoids commonly develop complex internal architecture and central necrotic cores^18^, quantitative assessments of viability and differentiation vary substantially with sectioning depth. We illustrate this using multiplexed immunofluorescence of mouse whole-brain organoids to map developmental trajectories under static and shaken culture conditions at days 7, 14, and 21 (**Figure 2a-f**). CORE captured the full heterogeneous cohort at its true equatorial plane within a single section that exposed the fully developed internal architecture (section 60; **Figure 2a**). Quantification of total organoid footprint and necrotic core area, defined by the absence of neuronal MAP2 and astrocytic Aldh1l1 signal, revealed a systematic depth-dependent quantification bias (**Figure 2b, c; Supplementary Figure 3a**). In peripheral sections that bypass the necrotic core, marker-positive areas were inflated, artificially overstating the apparent differentiation state, while the coefficient of variation was more than twofold higher than at the equatorial plane (**Figure 2d**). These results demonstrate that section depth can substantially influence quantitative readouts, potentially obscuring biologically meaningful differences between experimental conditions.

**Figure 2.**
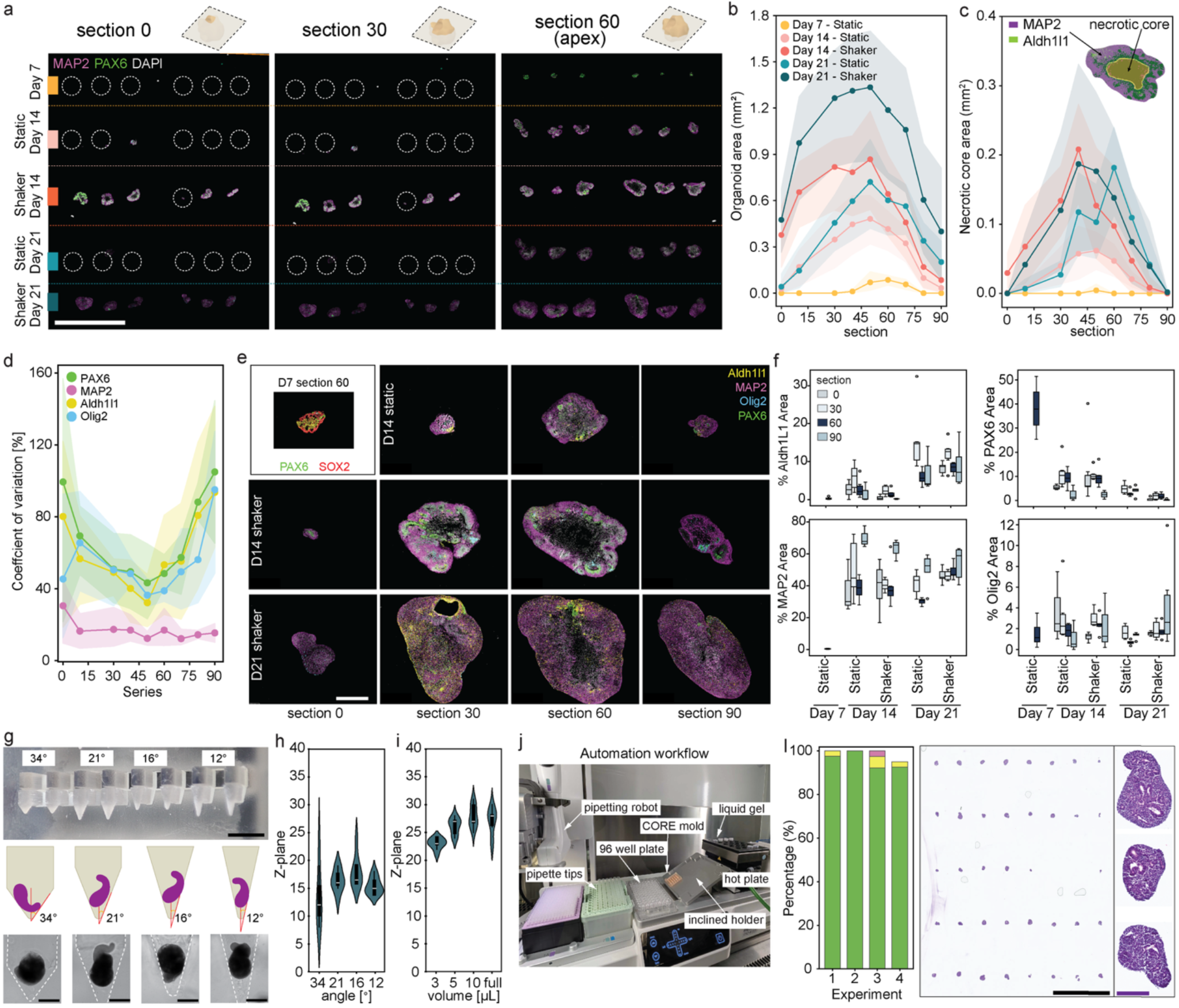
**CORE enables unbiased biological profiling and automation across 3D tissue models**. **a**, Multiplex immunofluorescence (mIF) across three sections of a 5×6 CORE array, showing coplanar alignment and capture of the full organoid cohort at the equatorial place (section 60). Scale bar, 5 mm. **b, c**, Total organoid area (**b**) and necrotic core area (**c**) as a function of section depth, revealing depth-dependent sampling vias in quantitative measurements (necrotic regions defined by MAP2−/Aldh1l1− signal loss; shaded areas: s.d., n ≥ 3). **d**, Marker coefficient of variation (CV) across conditions, demonstrating reduced measurement variability at the equatorial plane. **e**, Representative IF images across developmental stages (days 7, 14, 21) showing SOX2+ rosettes and localized PAX6 expression. Scale bar, 0.5 mm. **f**, Temporal marker dynamics in static and shaken conditions. Box plots demonstrate that accelerated neural maturation in shaken cultures is detectable at the equatorial plane but becomes obscured in peripheral sections. **g**, Schematic and photographs of CORE inserts with cone apertures ranging from 33.7° to 12°; brightfield images illustrate improper wall-resting of gastruloids at 33.7°. Scale bar, 0.5 mm. **h, i**, Gastruloid axial position distributions across cone apertures (**h**) and anchoring hydrogel volumes (**i**) (n = 8). **j**, Integrated workstation featuring the ASSIST PLUS liquid-handling platform. **k**, Gastruloid alignment relative to the cone axis across four automated runs (n = 4). **L)** Representative H&E section from an automatically embedded gastruloid block demonstrating preservation of tissue architecture across the cohort. Scale bars, 5 mm (black) and 250 µm (purple).

Tracking these cohorts over a three-week culture timeline revealed how structural heterogeneity and variance emerge at day 14 and become pronounced by day 21 (**Figure 2e**). By capturing the central core, equatorial sampling encompasses the full radial architecture of the tissue and provides a more representative measure of whole-organoid composition. Using this unbiased sampling plane, CORE revealed accelerated neural maturation in shaken cultures at day 21, a difference detectable at the equatorial section (**Figure 2f**, section 60). In contrast, peripheral sections bypass the core and overestimate marker-positive areas, reducing the apparent separation between culture conditions and masking biological divergence. Collectively, these findings demonstrate that section-depth sampling bias can influence biological interpretation and highlight the importance of deterministic specimen alignment for reproducible tissue profiling.

While CORE excels with symmetric organoids, small, asymmetric models such as gastruloids present additional challenges^19^. Standard histology frequently results in sample loss and unreliable inter-sample comparison owing to imprecise embedding and axial misorientation. Consequently, researchers often rely on whole-mount 3D imaging^20^ despite labor-intensive clearing and staining workflows, variable antibody penetration, and limited side-by-side comparability of experimental conditions. To define the operating range of CORE for such specimens, we evaluated four cone aperture angles using mouse gastruloids, which possess a pronounced anteroposterior axis (**Figure 2g**). Narrower apertures reduced axial variance, with the 12.0° geometry producing the tightest distribution, likely owing to reduced hydrodynamic drag during backfilling (**Figure 2g, h; Supplementary Figure 3b**). During hydrogel encapsulation, we observed intermittently displacement of specimens from the apex, which we attribute to hydrodynamic drag and buoyant displacement during backfilling. A localized 3 µL anchoring step before bulk encapsulation suppressed this effect, whereas larger volumes increased positioning error (**Figure 2i, Supplementary Figure 3c**). Unlike conventional flat histomolds arrays^2^, CORE’s passive geometry-driven alignment readily integrates with automated liquid handling hardware. Using the optimized 12.0° geometry, we coupled CORE to a robotic liquid handler with an inclined holder and achieved automated end-to-end specimen transfer from 96-well plates to embedded blocks (**Figure 2j, Supplementary Figure 3d; Supplementary Video 2**). The automated pipeline achieved a 95.5% apex trapping efficiency across 160 total specimens (n=40 per run, four independent trials), (**Figure 2l**), yielding sections that captured internal tissue architecture across the entire gastruloid cohort.

In summary, CORE converts stochastic, depth-biased histopathology into a deterministic, geometry-driven workflow. By replacing manual specimen positioning with passive coplanar alignment, CORE eliminates spatial sampling variance, enables comparable single-section readouts across heterogeneous specimens, and recovers biological differences that depth artefacts would otherwise mask. As organoid models become increasingly important for screening, disease modeling, and spatial omics, removing this source of bias supports reproducible comparison across conditions, timepoints, and laboratories.

## Online Methods

### 3D design and fabrication of CORE parts

The three-dimensional architecture of the parts was designed using computer aided design software (Fusion 360, Autodesk). The resulting STL files (available as Supplementary Files) were imported into PreForm (Formlabs) and printed using a stereolithography system (Form 3B+, Formlabs). To ensure maximal planarity of the base, the bottom surface was printed directly on the printer bed without support. Spacer and Cone inserted were fabricated using High Temp resin (Formlabs), with a layer height of 25 µm. Following printing, all parts were washed and cured according to manufacturer’s protocols: 20-min bath in isopropanol under flow (Form Wash, Formlabs) followed by a 30-min UV-curing at 60 °C (Form Cure, Formlabs).

### CORE hydrogel mold preparation

The specific mold geometry was selected based on the dimensions and panel size of organoids to be processed. HistoGel™ (Epredia, referred to as ‘gel’ in this manuscript) was thawed in a microwave for ∼10-20 s and maintained in a liquid state at ∼ 64ºC in a bath throughout the workflow. Upon assembly of the mold, Histogel was pipetted into the open corner ensuring the absence of air injection. The inherent hydrophobicity of the resin surface allowed for complete gel filling without overflowing at the Cone Insert-Base interface. To facilitate the evacuation of any trapped air, the mold was slightly tilted towards the opposite side of the injection point. To account for gel contraction during polymerization, an excess of gel was applied at the injection port to prevent the entrapment of air. The Histogel was then polymerized at 4ºC for 20 min in a fridge. Following polymerization, the Spacer was removed by pulling vertically, while the Cone insert was removed by a sequential lateral shift and a vertical pull. The resulting CORE Histogel mold was either retained within the base for organoid loading or gently demolded using a spatula to squeeze it out and transferred to a petri dish or Parafilm for subsequent processing.

### Sample transfer and positioning

Biological samples were transferred using truncated P1000 pipette tips (Eppendorf) pre-coated with 1% BSA (130-091-376, MACS) in PBS to accommodate varying specimen sizes; alternatively, spatulas or tweezers were employed for manual manipulation. Specimens were aspirated with excess liquid (10-20 µL), allowed to sediment within the tip, and subsequently dispensed into individual wells using as little volume as possible. In cases where wells were pre-filled with liquid, the pipette tip was placed in direct contact with the well to facilitate gravitational sedimentation. To facilitate sample identification after processing, specimens were embedded in an asymmetric layout. A black support base, angled at 60°, was used to promote sedimentation into the cone apices while providing visual contrast for microscopy imaging. Excess liquid was removed by placing a folded precision wipe (Kimtech Science) atop the Histogel mold to absorb fluid via capillary action while retaining samples within the cones. This dehydration step can be repeated to further assist in positioning samples at the cone tips. Approximately 3 µL of liquid Histogel was added to fix each sample, and after 2 min, the wells were filled entirely, and the resulting Histogel block was placed horizontally at 4°C for 20 min. Upon removal from the base, the top right corner was excised. The histoarray was then placed in a histology cassette where a fragment of 4% PFA-fixed chicken breast tissue (stored in PBS with sodium azide at 4°C) was added at the corner as a position reference to be processed alongside the samples. Consequently, each stained section contains a section of chicken breast at the top-right corner. Finally, the cassette was immersed in 10% formalin for 4 h at room temperature.

### Paraffin embedding and sectioning

Following fixation, Histogel blocks were processed in a fully automated HistoCore PEARL tissue processor (Leica Biosystems). The protocol initiated with a 5-min delayed start in formalin at 35 °C with the mixing option off, followed by a graded dehydration series: two 30-min baths in 70% ethanol, one 60-min bath in 80% ethanol, two 30-min baths in 96% ethanol, and two 60-min baths in 100% ethanol. Clearance was achieved through two 45-min xylene immersions, followed by paraffin infiltration across three separate baths at 56°C for 45 mins each. Subsequent embedding was performed on a Medite TES99 station (MEDITE Medical GmbH). Processed Histogel blocks were transferred to a metallic mold without flipping; the block was gently pressed against the mold base on a cold plate to ensure the cast surface was coplanar with the cutting plane before the mold was filled with liquid paraffin and solidified. Sectioning was conducted on a microtome (Thermo Microm HM355S, Fisher Scientific); blocks were trimmed at 10–20 µm and cooled in an ice bath for 10–15 min to facilitate ribbon formation prior to cutting 3–4 µm sections. Sections were floated on a distilled water bath, maintained at 42 °C and subsequently mounted on Superfrost or Gold Superfrost slides for overnight curing at 37 °C.

### Hematoxylin and Eosin (H&E) Staining

Automated H&E staining was executed following the standard protocol on the Ventana HE600 stainer (Roche Tissue Diagnostics).

### Multiplexed immunofluorescence

Opal fluorophores were prepared according to the manufacturer’s instructions. All primary antibodies used for immunofluorescence and multiplex staining are detailed in **Supplementary Table 1**. Multiplexed staining was performed using a Ventana Discovery Ultra automated tissue stainer (Roche Tissue Diagnostics). Validated primary antibodies were diluted in Discovery Ab diluent, with the specific application order established during preliminary runs. The automated protocol commenced with three cycles of baking and deparaffinization (60°C for 8 min, followed by 69°C for 8 min). Then, heat-induced antigen retrieval was conducted using Tris-EDTA buffer (pH 7.8) at 95°C for 40 min, followed by blocking with Discovery Goat Ig Block for 32 min. Subsequent staining cycles involved the sequential application of Discovery Inhibitor (16 min), primary antibody (40 min), species-specific HRP-conjugated secondary antibody (16 min), and the corresponding Opal dye (12 min at 37°C). To prevent cross-reactivity, an antibody neutralization and heat-denaturation step (92°C for 4 min) was performed between each cycle. For the Opal 780 sequence specifically, an additional Goat IgG blocking step (16 min) was performed after the primary antibody, followed by the secondary antibody-HRP, Opal TSA reagent, and Opal dye 780. Finally, slides were manually mounted using ProLong™ Gold Antifade Mountant (Invitrogen, Thermo Fisher Scientific) and allowed to dry for at least 2 hours prior to imaging.

### Organoid generation

Mouse iPSC-derived retinal organoids were generated and previously described by Völkner et al^21^. Mouse whole-brain organoids were generated by adapting a protocol for human unguided cerebral organoids^22^ to mouse embryonic stem cells. Mouse embryonic stem cell-derived gastruloids were produced following the detailed protocol described by Vianello et al^23^.

### Confocal microscopy

Following embedding, Histogel blocks were permeabilized in 1× PBS supplemented with 0.5% Triton X-100 (Sigma-Aldrich) for 2 h at room temperature. Immunostaining was conducted in a blocking buffer (2% normal donkey serum and 0.05% Triton X-100 in PBS). Blocks were incubated for 2 h with DAPI (1:1000; Invitrogen) and Alexa Fluor™ Plus 647 Phalloidin (1:2000; Thermo Fisher Scientific), followed by three 30-min washes in PBS. Images were acquired using an inverted microscope (Ti2, Nikon) equipped with a spinning disk confocal system (CSU-W1, Yokogawa) and a Orca-Fusion sCMOS camera (C14440-20UP, Hamamatsu). Imaging was performed using a Plan Fluor 10x objective (NA 0.3). Z-stacks consisting of 59 optical sections were collected with a 25 µm step size. Fluorescence excitation was achieved using 405 nm and 561 nm laser lines for DAPI and Cy3 channels, respectively, with exposure times set to 300 ms. Image acquisition and system control were managed using Nikon NIS-Elements AR software (version 5.42.05).

### Automated organoid transfer

Automated organoid transfer was performed using the ASSIST PLUS pipetting robot (Integra Biosciences) equipped with a D-ONE single-channel pipetting module (0.5–300 µL range). To minimize sample adhesion, wide-bore pipette tips were pre-coated with 1% BSA (130-091-376, MACS) in PBS via three mixing cycles (50 µL). During the transfer procedure, the receiving Histogel mold was positioned at a 30° inclination. Organoids were captured from U-bottom source plates by aspirating 10 µL (speed level 5) at a height of 0.3 mm from the well bottom. Samples were dispensed into the upper region of the Histogel mold wells at a low flow rate (speed level 1) with a horizontal x-offset of 0.6 mm. Following transfer, the mold was tilted to 90° and gently agitated to facilitate organoid sedimentation before returning to the 30° inclination. Excess medium (5 µL) was removed using standard pipette tips at minimum speed (speed level 1). Encapsulation was completed by dispensing 10 µL of Histogel into the center of each well using wide-bore tips (speed level 3). Molds were subsequently imaged and processed for paraffin embedding as described above.

### Histology analysis

Organoid boundary areas were traced based on raw DAPI and structural marker signals. The localized internal necrotic core area was explicitly segmented and quantified by mapping the internal regions presenting a total absence of MAP2 and Aldh1l1 fluorescence HALO AI software (Indica Labs, v3.6.4134). Individual marker coverage thresholds were calculated as percentage areas relative to the total segmented organoid footprint. The coefficient of variation (CV) across depth tracks was computed using standard analytical formulations across aligned biological replicates (n >= 3). Data visualization, boxplots, and statistical verification were compiled using custom Python scripts.

## Acknowledgments

We thank G. Brancati for providing the retinal organoids used in this study; S. Tsiepa for technical assistance with tissue sectioning and H&E staining; R. Hans, M. Girgin and G. Rossi for establishing the gastruloid protocols; and L. Pollich for producing the CORE mold demonstration video.

## Author Contributions

I.P. conceived the project, designed and performed experiments, prepared the figures, and wrote the original manuscript draft. B.L. contributed to the conceptualization of the study, designed and performed experiments, assisted with figure preparation, and contributed to manuscript writing and editing. C.L.-B. and S.T. performed experiments. J.A. contributed to the conceptualization of the study and the initial proof of concept. I.C. performed histological processing. J.R. provided the adapted brain organoid protocol, established antibody panels, and contributed to scientific discussions. J.L.G.-C. contributed to the conceptualization of the study, provided scientific oversight, contributed to figure preparation and revision, guided data interpretation, and contributed to manuscript writing, reviewing, and editing.

---

**Supplementary Video 1:** Disassembly of CORE mold.

**Supplementary Video 2:** Automated organoid transfer to the CORE mold.

**Supplementary Figure 1:**
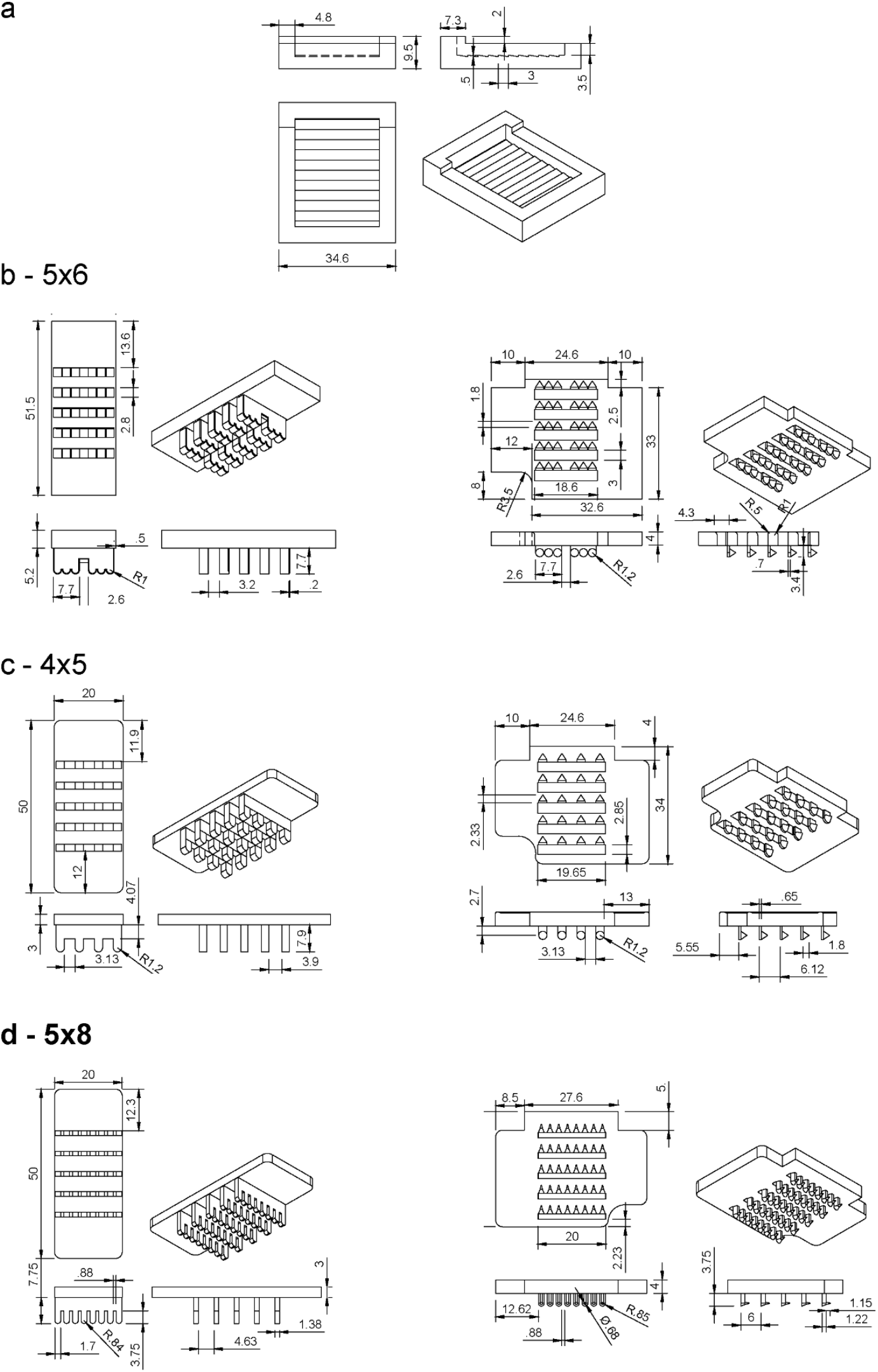
**a**, Technical drawings with dimensions of generic base part, compatible with all spacer and cone insert arrays. (**b**: 5×6, **c**: 4×5, **d**: 5×8) Three different array designs featuring their respective Spacers, left, and Cone Inserts, right.

**Supplementary Figure 2:**
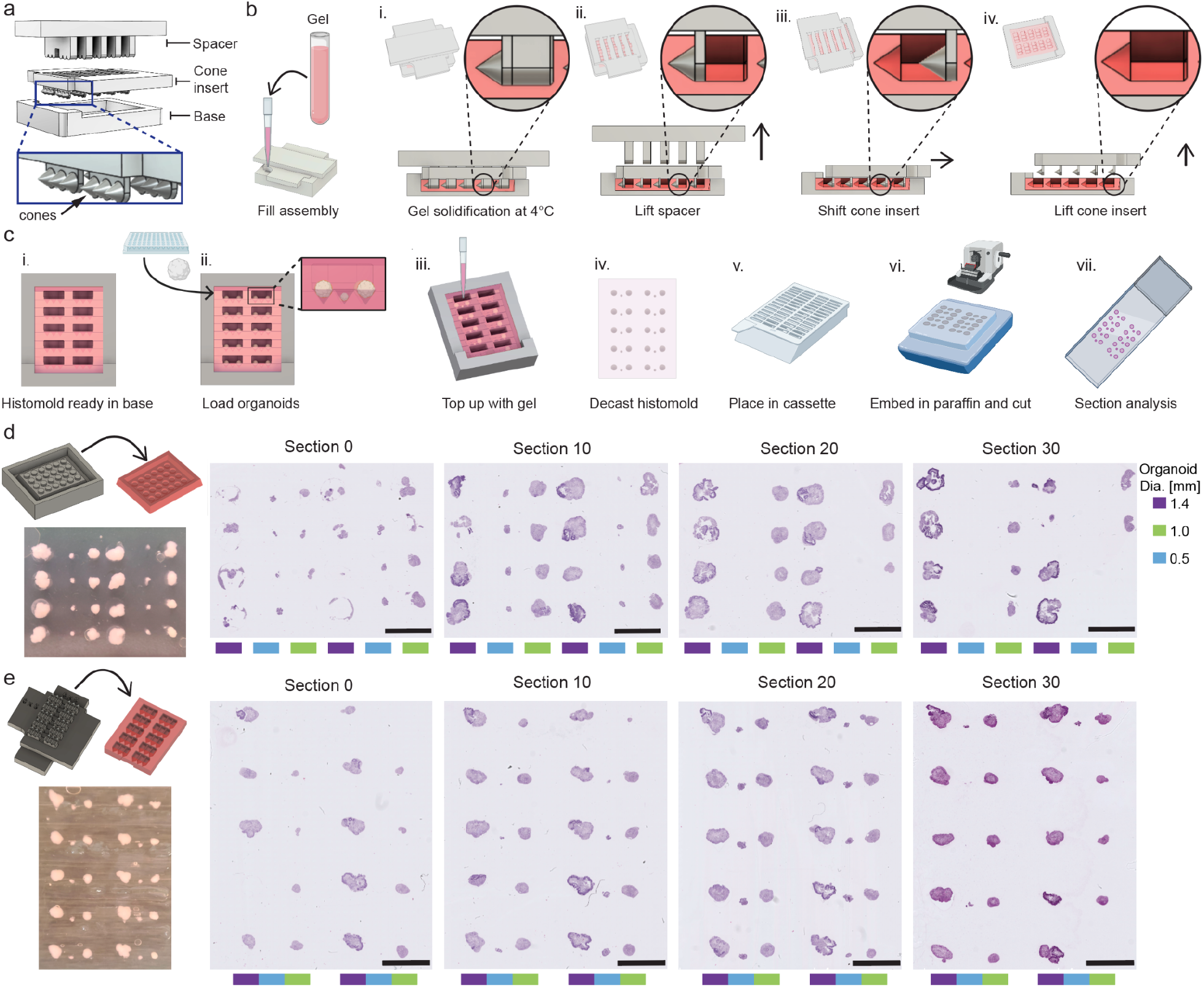
**a**, Exploded view of the three-part mold assembly: Base, Spacer, and Cone Insert. **b**, Schematic of the demolding kinetics showing the shift-to-release strategy: (i) gel solidification, (ii) vertical removal of the Spacer, (iii) lateral displacement of the Cone Insert, and (iv) vertical withdrawal to preserve the delicate gel features. **c**, Schematic of the operational workflow: (i) gel casting, (ii), organoid loading, (iii) gravity-assisted alignment and gel anchoring, (iv) decasting, (v) fixation in cassette, (vi) paraffin embedding and microtome sectioning, and (vii) slide mounting and staining. **d, e**, Schematic and H&E sections of retinal organoids embedded in a flat-bottomed hydrogel mold (**d**) and the CORE platform (**e**), across serial sections obtained at 5 µm intervals. Scale bars, 5 mm.

**Supplementary Figure 3:**
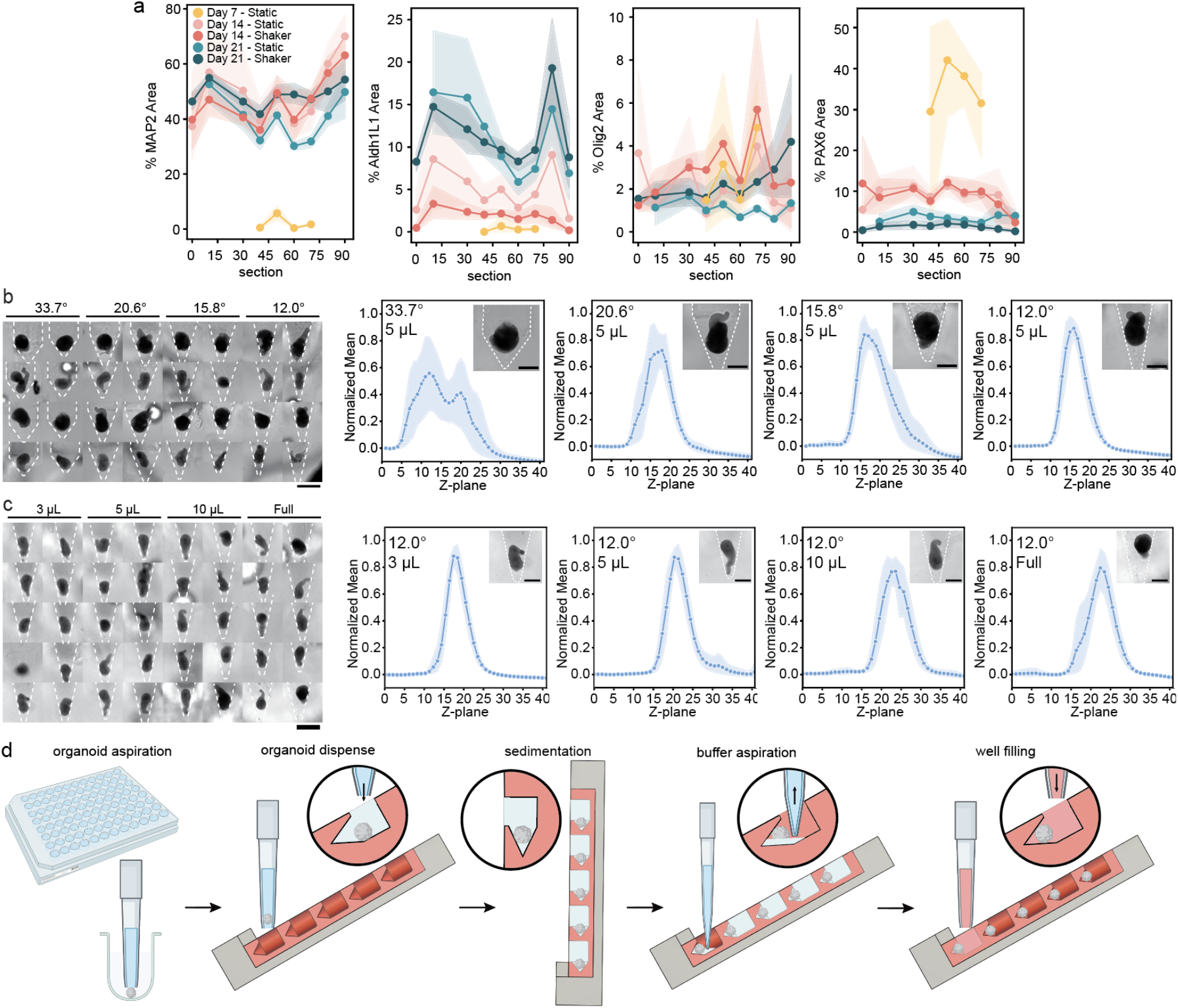
**a**, Marker quantification expressed as a percentage of total organoid area across serial sections to track neurons, astrocytes, and oligodendrocytes (Olig2). **b**, On the left, representative bright-field images of mouse gastruloids embedded within different apertures. Scale bar, 1 mm. On the right, normalized fluorescence distribution profiles across the z-axis, showing the narrowest axial distribution in the 12° aperture. Shaded areas indicate standard deviation (n=8 organoids). Scale bar, 0.5 mm. **c**, Bright-field images illustrating the effect of initial pre-fixing volumes (3 μL, 5 μL, 10 μL, and Full) on gastruloid positioning. Scale bar, 1 mm. Normalized fluorescence distribution profiles across the z-axis confirming that a 3 μL prefixing volume yields the most symmetrical and reproducible coplanarity. Shaded areas indicate standard deviation (n=10 organoids). Scale bar, 0.5 mm. **d**, Schematic of the automated organoid embedding workflow, illustrating the sequential steps: dispensing of specimens, sedimentation (at 90º), aspiration of buffer, and embedding with supplementary gel. Schematic was partially created with BioRender.com.

**Supplementary Table 1.** List of primary antibodies used for immunofluorescence and multiplex staining.

|  | **Antibody** | **Species** | **Clonality** | **Dilution** | **Company** | **Catalog number** |
| --- | --- | --- | --- | --- | --- | --- |
| 1 | Pax6 | Rabbit | Polyclonal | 1/500 | Biolegend | 901301 |
| 2 | MAP2 | Mouse | Monoclonal | 1/100 | Merck | MAB3418 |
| 3 | ALDH1L1 | Mouse | Monoclonal | 1/100 | Millipore | MABN495 |
| 4 | Olig2 | Rabbit | Monoclonal | 1/100 | Abcam | ab109186 |
| 5 | SOX2 | Goat | Polyclonal | 1/100 | R&D | AF2018 |

## References

1. Heub, S. et al. Coplanar embedding of multiple 3D cell models in hydrogel towards high-throughput micro-histology. Sci. Rep. 12, 9991 (2022).

2. Harter, M. F. et al. High-throughput histopathology for complex in vitro models. Cell Rep. Methods 6, 101419 (2026).

3. Olsen, Timothy. R. et al. Processing cellular spheroids for histological examination. J. Histotechnol. 37, 138–142 (2014).

4. Vuille-dit-Bille, E. et al. An acoustic levitation platform for high-content histological analysis of 3D tissue culture. Lab a Chip 25, 2732–2743 (2025).

5. Zhang, S. et al. An efficient and user-friendly method for cytohistological analysis of organoids. J. Tissue Eng. Regen. Med. 15, 1012–1022 (2021).

6. Weisler, W., Miller, S., Jernigan, S., Buckner, G. & Bryant, M. Design and testing of a centrifugal fluidic device for populating microarrays of spheroid cancer cell cultures. J. Biol. Eng. 14, 7 (2020).

7. Yoshikawa, A. L., Omura, T., Takahashi-Kanemitsu, A. & Susaki, E. A. Blueprints from plane to space: outlook of next-generation three-dimensional histopathology. Cancer Sci. 115, 1029–1038 (2024).

8. Lee, J. et al. Adaptive compression framework for giga-pixel whole slide images. Nat. Commun. 17, 207 (2025).

9. Wang, K. et al. Toward universal immunofluorescence normalization for multiplex tissue imaging with UniFORM. Cell Rep. Methods 5, 101172 (2025).

10. Sater, S. et al. Society for Immunotherapy of Cancer: Standards for Reporting of Multiplex Immunohistochemistry/Immunofluorescence Assays (STORMI). J. Immunother. Cancer 13, e012280 (2025).

11. Koh, I. & Hagiwara, M. Gradient to sectioning CUBE workflow for the generation and imaging of organoids with localized differentiation. Commun. Biol. 6, 299 (2023).

12. Yang, C. et al. 4D-Printed Transformable Tube Array for High-Throughput 3D Cell Culture and Histology. Adv. Mater. 32, e2004285–e2004285 (2020).

13. Vuille-dit-Bille, E. et al. PEGDA-based HistoBrick for increasing throughput of cryosectioning and immunohistochemistry in organoid and small tissue studies. Sci. Rep. 15, 412 (2025).

14. Gabriel, J., Brennan, D., Elisseeff, J. H. & Beachley, V. Microarray Embedding/Sectioning for Parallel Analysis of 3D Cell Spheroids. Sci. Rep. 9, 16287 (2019).

15. Kang, J. et al. Mini-pillar array for hydrogel-supported 3D culture and high-content histologic analysis of human tumor spheroids. Lab a Chip 16, 2265–2276 (2016).

16. Chen, J. et al. Cerebral Organoid Arrays for Batch Phenotypic Analysis in Sections and Three Dimensions. Int. J. Mol. Sci. 24, 13903 (2023).

17. Biunno, I., Paiola, E. & Blasio, P. D. The Application of the Tissue Microarray (TMA) Technology to Analyze Cerebral Organoids. J Histochem Cytochem. 69, 451–460 (2021).

18. Lancaster, M. A. & Knoblich, J. A. Generation of cerebral organoids from human pluripotent stem cells. Nat. Protoc. 9, 2329–2340 (2014).

19. Rossi, G., Giger, S., Hübscher, T. & Lutolf, M. P. Gastruloids as in vitro models of embryonic blood development with spatial and temporal resolution. Sci. Rep. 12, 13380 (2022).

20. Rossi, G. et al. Capturing Cardiogenesis in Gastruloids. Cell Stem Cell 28, 230–240.e6 (2021).

21. Völkner, M. et al. Retinal Organoids from Pluripotent Stem Cells Efficiently Recapitulate Retinogenesis. Stem Cell Rep. 6, 525–538 (2016).

22. Giandomenico, S. L., Sutcliffe, M. & Lancaster, M. A. Generation and long-term culture of advanced cerebral organoids for studying later stages of neural development. Nat. Protoc. 16, 579–602 (2021).

23. Vianello, S. D., Girgin, M., Rossi, G. & Lutolf, M. Protocol to generate Gastruloids (LSCB, EPFL) v1. protocols.io 10.17504/protocols.io.9j5h4q6 (2019) doi:10.17504/protocols.io.9j5h4q6.

